# In vivo relevance of intercellular calcium signaling in *Drosophila* wing development

**DOI:** 10.1101/187401

**Authors:** Qinfeng Wu, Pavel A. Brodskiy, Francisco Huizar, Jamison J. Jangula, Cody Narciso, Megan Levis, Teresa Brito-Robinson, Jeremiah J. Zartman

**Affiliations:** Department of Chemical and Biomolecular Engineering, University of Notre Dame, 205 McCourtney Hall, Notre Dame, IN 46556, USA

**Keywords:** calcium wave, quantitative biology, signal transduction, dynamic systems, mechanobiology

## Abstract

Recently, organ-scale intercellular Ca^2+^ transients (ICTs) were reported in the Drosophila wing disc. However, the functional in vivo significance of ICTs remains largely unknown. Here we demonstrate the in vivo relevance of intercellular Ca^2+^ signaling and its impact on wing development. We report that Ca^2+^ signaling in vivo decreases as wing discs mature. Ca^2+^ signaling ex vivo responds to fly extract in a dose-dependent manner. This suggests ICTs occur in vivo due to chemical stimulus that varies in concentration during development. RNAi mediated inhibition of genes required for ICTs results in defects in the size, shape, and vein patterning of adult wings. It also leads to reduction or elimination of in vivo Ca^2+^ transients. Further, perturbations to the extracellular matrix along the basal side of the wing disc stimulates intercellular Ca^2+^ waves. This is the first identified chemically defined, non-wounding stimulus of ICTs. Together, these results point toward specific in vivo functions of intercellular Ca^2+^ signaling to mediate mechanical stress dissipation and ensure robust patterning during development.

## Introduction

Calcium (Ca^2+^) signaling provides a versatile and highly conserved toolkit for modulating cellular mechanics, cellular differentiation, proliferation, wound healing and regeneration during animal development (Prevarskaya et al., 2011; Markova and Lenne, 2012; Monteith et al., 2012; Antunes et al., 2013; Deng et al., 2015; Restrepo and Basler, 2016; Wallingford et al., 2001). Cells encode complex signals through the amplitude and frequency of oscillations of cytoplasmic Ca^2+^ concentrations. Cells decode these signals by modulating the activities of downstream enzymes and transcription factors through reversible binding of Ca^2+^ to Ca^2+-^binding domains (Berridge et al., 2000; Clapham, 2007; Smedler and Uhlén, 2014). Although extensively studied in neuronal and cardiac cells, Ca^2+^ signaling also occurs in epithelial tissues. However, its specific roles in epithelial development are poorly understood.

Regulation of cytoplasmic Ca^2+^ levels can occur through controlled release of Ca^2+^ from the endoplasmic reticulum (ER). Inositol trisphosphate (IP_3_) is a small molecule that binds to IP_3_ receptor (IP_3_R, Fig. 1A), which is a highly-conserved channel regulating Ca^2+^ release from the ER (Agrawal and Hasan, 2012; Clapham, 1995). Sarco/endoplasmic reticulum Ca^2+^-ATPase (SERCA) pumps Ca^2+^ back into the ER to replenish Ca^2+^ storage (Mekahli et al., 2011) and reduces the toxicity of long-term elevated Ca^2+^ concentrations (Berridge et al., 2000). Phospholipase C (PLC) is a family of catalysts that produces IP_3_ (Shortridge and McKay, 1995). Both Ca^2+^ and IP_3_ diffuse through gap junctions to propagate signals (Leybaert and Sanderson, 2012). The store-operated channels (SOCs), Stim and Orai, refill the ER (Feske et al., 2006; Roos et al., 2005).

**Fig. 1.**
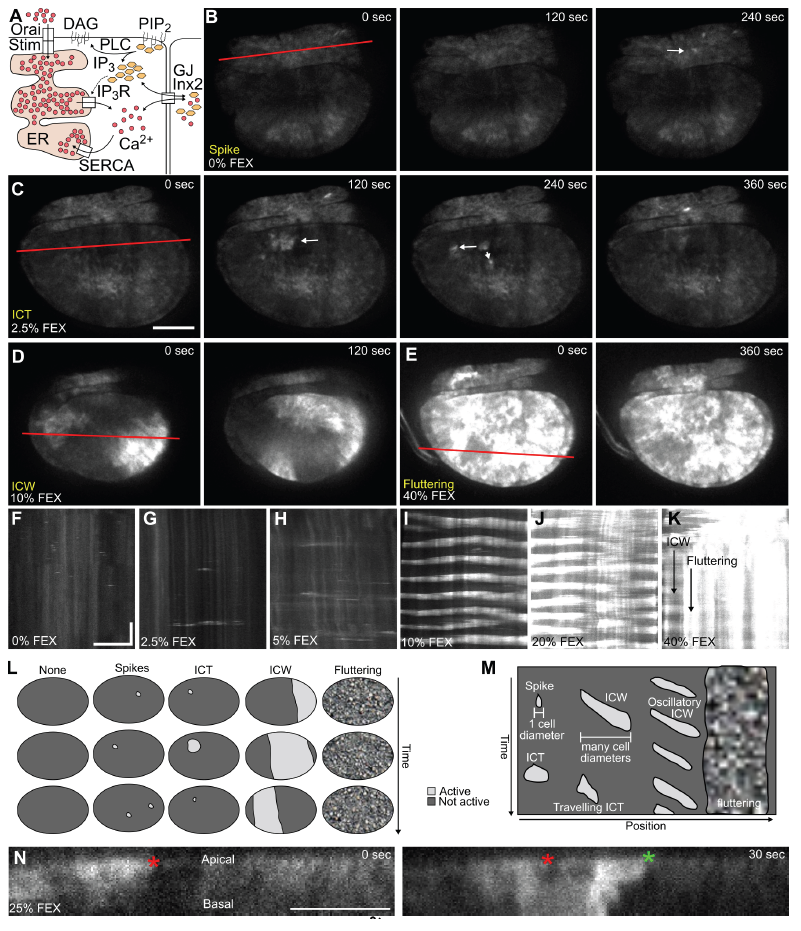
ICTs ex vivo exhibits a range of Ca ^2+^ activities in a dose-dependent manner.

A) Schematics of key Ca^2+^ signaling pathway components. IP_3_: Inositol trisphosphate; IP_3_R: IP_3_ receptor; gap junction: Gap junctions; Inx2: Innexin 2; ER: endoplasmic reticulum; SERCA: Sarco/endoplasmic reticulum Ca^2+^-ATPase; PIP_2_: phosphatidylinositol 4,5-bisphosphate; DAG: diacylglycerol. B-E) Live imaging of wing discs expressing UAS-GCaMP6f under nub-Gal4 in chemically defined ZB media with varying concentrations of fly extract. B) 0%, C) 2.5%, D) 10%, E) 40% v/v fly extract. These conditions stimulate four Ca^2+^ activities: spikes (0%), ICTs (2.5%), ICWs (10% v/v fly extract) and “fluttering” Ca^2+^ oscillations (40% v/v fly extract). F-K) Kymographs parallel to the dorsal/ventral axis of the pouch under different concentrations of fly extract: F) 0%, G) 2.5%, H) 5%, I) 10%, J) 20%, K) 40%. Scale bars represent 50 μm and 10 minutes. L-M) Cartoon of four main Ca^2+^ activities. L) Timelapse cartoons showing spikes, ICTs, ICWs, and fluttering. M) Kymograph cartoon showing spikes do not propagate to neighboring cells, ICTs propagate a short distance, ICWs propagate many cell diameters, ICWs may be single events or oscillatory, and fluttering is a long-term state. N) Orthogonal sections through a wing disc undergoing a calcium transient reveals 3D anisotropy in Ca^2+^ propagation. Red and green asterisks indicate wave front at time 0 s and 30 s respectively. White arrows indicate Ca^2+^ oscillations. Scale bars represent 50 μm.

Recent results from our group (Narciso et al., 2017) and others (Balaji et al., 2017; Restrepo and Basler, 2016) suggest that intercellular Ca^2+^ transients (ICTs) occur spontaneously in wing discs in vivo and that ICTs can also be stimulated by fly extract ex vivo. Although previously reported, the in vivo relevance and significance of ICTs has been questioned, largely because a systematic analysis has not been performed. Previous studies and an accompanying report (Brodskiy et al, DEVELOP/2017/159269) focus primarily on Ca^2+^ dynamics in ex vivo cultures because in vivo imaging suffers from larval motion, mechanical compression and inability to specifically perturb wing discs in vivo with drugs. Consequently, the dynamics and functional roles of ICTs during development are poorly understood.

Here, we demonstrate that Ca^2+^ signaling regulates tissue growth, vein patterning and morphogenesis of the wing disc. Specific defects include smaller wings, disappearance of veins and ectopic folding of the wing disc. Qualitatively, both ex vivo and in vivo imaging of Ca^2+^ share exhibit the same four categories of Ca^2+^ activities in wing discs. Strikingly, the release of tissue stress by the dissociation of extracellular matrix (ECM) alone induces ICTs in wing disc ex vivo culture without fly extract. This represents the first non-wounding, chemically defined perturbation that stimulates ICTs. This stimulation of ICTs by removing the mechanical constraint suggests a specific function of ICTs. In particular, stochastic in vivo ICT dynamics mediate remodeling and expansion of the tissue during tissue growth. We propose that this “mechanical stress dissipation” hypothesis explains a large portion of intercellular calcium dynamics during development of growing organs.

## Results

### Oscillatory Ca ^2+^ signal exhibits four categories of Ca ^2+^ activities in wing discs

We began by systematically comparing the dynamics of ICTs, both ex vivo and in vivo, to understand the function of ICTs in wing development. We imaged larvae expressing nub-Gal4, UAS-GCaMP6 to investigate the Ca^2+^ dynamics in *Drosophila* wing discs. GCaMP6 is a genetically-encoded sensor that visualizes Ca^2+^ through fluorescence (Sun et al., 2013). We incubated wing discs with different concentrations of fly extract and performed live imaging of Ca^2+^ signaling that produce a range of ICTs ex vivo. We discerned four categories of cytoplasmic Ca^2+^ oscillations in wing discs: single cell spikes (Fig. 1B, F), ICTs (Fig. 1C, G), periodic ICTs we term intercellular Ca^2+^ waves (ICWs, Fig. 1D, I, J), and an overstimulated phenotype of cytoplasmic Ca^2+^ oscillations we term “fluttering” (Fig. 1E, L, M). We discovered that in the absence of fly extract, only Ca^2+^ spikes occurred (Fig. 1B, F; Movie S1). Spikes are oscillations confined to individual cells, as observed in tissues like *Drosophila* guts (Deng et al., 2015) and *Xenopus* neural crests (Belgacem and Borodinsky, 2011). At low levels of fly extract (2.5%), stochastic, short-distance ICTs occur (Fig. 1C, G; Movie S2). Intermediate doses (5% to 20%) result in persistent ICWs (Fig. 1D, H-J; Movie S3-5). Higher doses of fly extract (40%) cause a loss of ICT activity and rapid Ca^2+^ oscillation due to continuous Ca^2+^ stimulation in cells (Fig. 1E, K; Movie S6). We also observed that Ca^2+^ signaling propagates preferentially along the apical surface than the basal surface (Fig. 1N). This is consistent with the higher concentration of gap junctions near the apical side of cells in imaginal tissues (Richard and Hoch, 2015). In this manuscript, we use ICTs to generally describe intercellular Ca^2+^ signaling activities encompassing spikes, ICTs and ICWs.

For in vivo ICTs, we gently immobilized the larvae with tape on a coverslip to image the ICTs.We observed the same four categories of ICTs during in vivo imaging: local spikes (Fig. 2A; Movie S7), ICTs (Fig. 2B; Movie S8), ICWs (Fig. 2C; Movie S9), and occasionally, fluttering discs (Fig. 2D; Movie S10). Overall, we observed ICTs in 40% of larva of different stages imaged during twenty-minute observation periods (n = 100). ICTs occurred more frequently in younger larvae than in older larvae (Fig. 2E, F). We conclude that ICTs are a recurring phenomenon in vivo in the absence of exogenous mechanical or chemical stimulation and correlate with developmental stages. It contradicts previous suggestions that ICTs may be an ex vivo artifact (Balaji et al., 2017; Dye et al., 2017).

**Figure 2:**
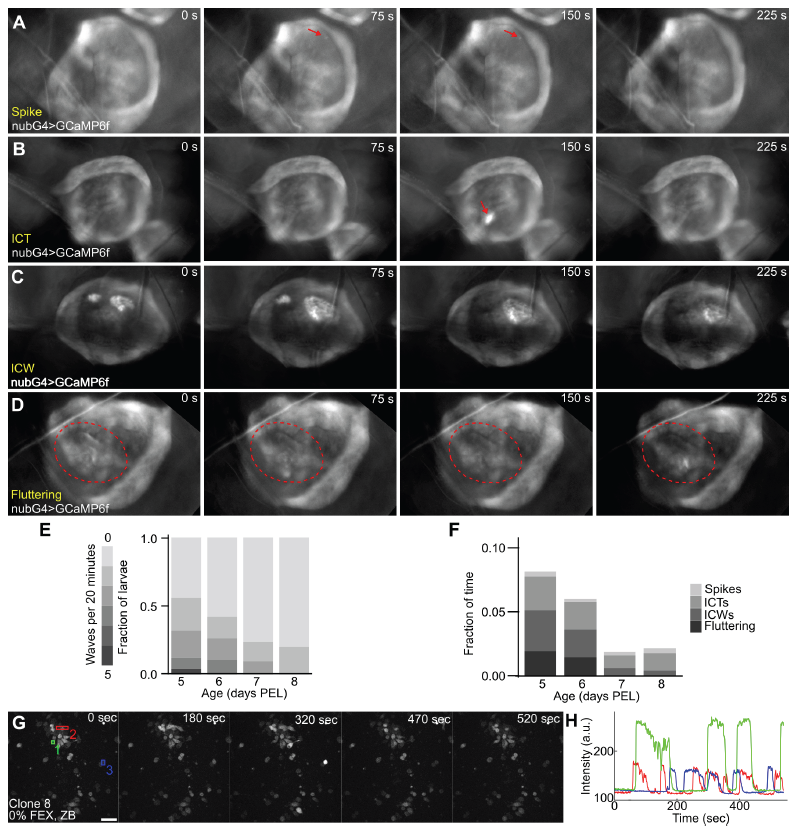
ICTs in vivo exhibits the same four different Ca ^2+^ activities as ex vivo wing discs and become less active with age. A-D) In vivo time-lapse fluorescent microscopy images of *nub*>*GCaMP6f* larval wing discs. A diverse range of ICT dynamics are observed, including: A) single-cell spike (red arrow), B) ICTs (red arrow), C) ICWs, and D) fluttering Ca^2+^ oscillation. The red circle indicates the area of fluttering Ca^2+^ oscillation. E) Number of ICT events encompassing all four categories were counted blindly in 20-minute long videos. Number of ICT events decreases with increasing larval age (p<0.018 by t-test of linear regression coefficient). F) The duration of each Ca^2+^ oscillation was counted blindly. The total Ca^2+^ activation time (the height of the column) decreases as wing discs grow older G) Ca^2+^ oscillations in Cl. 8 cells. H) Ca^2+^ dynamics of the colored cells showing oscillatory behaviors.

We also created a stable line of Clone 8 cells expressing GCaMP6 to further investigate the media requirements for ICTs in wing disc cells. Clone 8 cells were originally derived from cultured wing disc explants and are a useful tool for biochemical studies (Currie et al., 1988). Surprisingly, we observed Ca^2+^ oscillations in Clone 8 without fly extract (Fig. 2G, Movie S11) in a chemically defined media (Burnette et al., 2014). These oscillations occurred on the scale of roughly one per minute (Fig. 2H), which is an order of magnitude higher frequency than that observed in the wing discs stimulated with 15% fly extract. Thus, Ca^2+^ oscillations arise spontaneously in individual wing disc cells.

Cumulatively, these results indicate that the tissue microenvironment dampens Ca^2+^ signaling and that an active biochemical signaling in vivo and in fly extract stimulates coordinated intercellular Ca^2+^ oscillations. Ca^2+^ signaling as tissue level cell communication occurs through biochemical and mechanical cell-cell and cell-ECM signaling. Thus, stimulation of ICTs could be impacting these processes.

### The core Ca ^2+^ signaling toolkit is required for robust patterning, size and morphogenesis

We used RNA interference (RNAi) to knock down components of the Ca^2+^ signaling pathway to characterize the developmental functions of ICTs. We used multiple Gal4 drivers to express UAS-RNAi constructs either uniformly or differentially in the pouch to better define spatial and temporal requirements of Ca^2+^ signaling. Nub-Gal4 drives UAS-transgene expression uniformly in the 3^rd^ instar wing disc pouch (Fig. S1). MS1096-Gal4 has stronger expression in the dorsal than ventral compartments (Capdevila and Guerrero, 1994; Mirth et al., 2009). ap-Gal4 drives transgene expression exclusively in the dorsal compartment and is strongly expressed in earlier stages of development (2^nd^ instar) when the D/V boundary is specified (Hartl and Scott, 2014).

We inhibited genes known to be involved in Ca^2+^ signaling in the wing (Narciso et al., 2017). We used RyR^RNAi^ against ryanodine receptor (RyR) as a control. RyR is not expressed in the wing disc (Gramates et al., 2017). As expected, MS1096-Gal4 x UAS-RyR^RNAi^ did not exhibit morphological defects (Fig. 3A, B). We found that there are four primary wing defects in adult wings when perturbing Ca^2+^ signaling: 1) wing bending, 2) reduction in wing size, 3) loss of cross veins and 4) crumpled, blistered wings (Fig. 3C-H). Knocking down SERCA led to smaller, crumpled and blistered wings (Fig. 3C). Knocking down gap junctions through *MS1096*>*Inx2*^RNAi^ led to significantly smaller wings with defects including partial loss, total loss, and deviation in the posterior and anterior cross veins (PCV, ACV, Fig. 3D, G, H). We observed that partial loss of the PCV was always the loss of the posterior side of the PCV, and never the anterior side (n=33). *MS1096*>*IP_3_R*^RNAi^ exhibited similar but less severe wing defects (Fig. 3E, G), possibly because RNAi results in knockdown and not knockout of gene expression. Wing size was also significantly reduced (Fig. 3H). Overall, these results demonstrate that Ca^2+^ signaling plays important roles in wing morphogenesis, vein patterning, size control and Dpp secretion (Dahal et al., 2017, 20).

**Figure 3:**
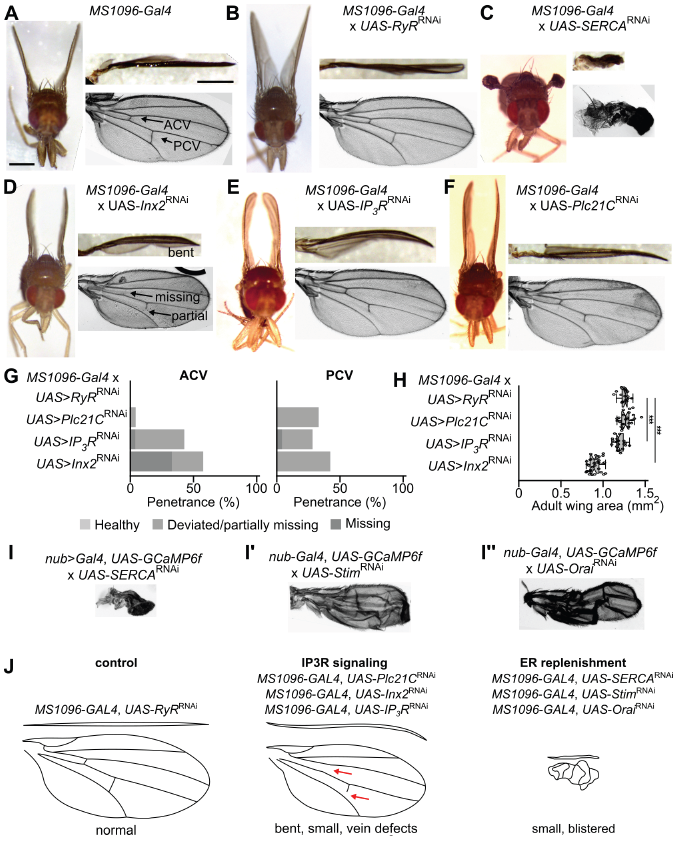
Perturbation of the intercellular Ca ^2+^ signaling impacts wing morphogenesis and vein patterning. A-F) Micrographs of adult fly, orthogonal view of wing, and mounted view of wing. A) Parental tester line (MS1096-GAL4). Arrows indicate anterior and posterior cross veins (ACV and PCV). B-F) MS1096-Gal4 line crossed with indicated RNAi lines (*UAS-RyR^RNAi^*; *UAS-SERCA^RNAi^*; *UAS-Inx2^RNAi^*; *UAS-IP_3_R^RNAi^*; *UAS-Plc21C^RNAi^*). Scale bars represent 0.5 mm. G) Fraction of wings with patterning defects in the ACV and PCV. Wings were scored as having a normal cross vein, a defective cross vein, or a missing cross vein. H) Total area of adult wings for the four indicated genotypes. I) Mounted wings from *nub*-*GAL4*, *UAS*-*GCaMP6f* crossed to UAS-*SERCA*^RNAi^ UAS-*Stim*^RNAi^ and UAS-*Orai*^RNAi^. J) Schematics of the different defects. *** indicates p<0.001 by Mann-Whitney *U* test. Sources of driver and UAS lines indicated in Table S1.

Three homologs of the PLC family are present in *Drosophila*: *Plc21C* (a PLCβ1 homolog), *sl* (*PLCγ*), and *norpA* (a PLCβ4 homolog). We observed that *MS1096*>*Plc21C*^RNAi^ wings had a similar bent phenotype and defects in the PCV to the *MS1096*>*IP_3_R*^RNAi^ and *MS1096*>*Inx2*^RNAi^ wings, but did not have the same ACV and wing size phenotypes (Fig. 3F, G). *nub*>*PLCγ*^RNAi^ had similar wings to controls but were smaller (Fig. S2), which has been reported to be caused by regulation of epidermal growth factor receptor signaling by the S2 domain of *PLCγ* which does not catalyze IP_3_ generation (Murillo-Maldonado et al., 2011). Overall, these results show that *Plc21C* is the primary PLC responsible for stimulating IP_3_R in the wing disc. This phenotype is thus linked to the observed inhibition of ICTs.

In contrast, we did not observe strong morphological defects when expressing *Plc21C* or *IP_3_R* RNAi under nub-Gal4 (Fig. S2). This difference may be due to the spatial difference of MS1096-Gal4 (higher expression in the dorsal compartment) and nub-Gal4 (uniform expression) drivers, which suggests that it is not only the inhibition of Ca^2+^ signaling, but also spatial heterogeneity in Ca^2+^ signaling that leads to folding in general.

Inhibition of *SERCA*, *Stim*, or *Orai* leads to chronic depression of Ca^2+^ concentrations in the ER. This induces apoptosis and reduces cell-cell adhesion (Mekahli et al., 2011). Smaller, blistered wings with thicker veins have been reported when *SERCA*, *Stim* or *Orai* is inhibited (Balaji et al., 2017; Eid et al., 2008; Restrepo and Basler, 2016). We confirmed that knocking down *SERCA*, *Stim*, and *Orai* led to smaller, blistered wings when driven by *nub* (Fig. 3I-I”).

These results indicate that inhibition of Ca^2+^ signaling components leads to two phenotypes depending on which part of the Ca^2+^ signaling toolkit is affected (Fig. 3J). Perturbation of the IP_3_ signaling and gap junction components (Plc21C, IP_3_R and Inx2) leads to smaller, bent wings with reduced cross veins. Perturbation of genes responsible for ER replenishment (SERCA, Stim and Orai) leads to blistered wings.

### The core Ca ^2+^ signaling is crucial for normal wing disc patterning and folding

To elucidate the mechanism of ICTs and its effect on wing disc development, we systematically knocked down components of the Ca^2+^ signaling pathway (Fig. 4A) and measured changes in ICT dynamics in vivo with crosses to the nub>GCaMP6 tester line. We use *nub>RyR^RNAi^* as the control (Movie S12). RNAi knockdown of *IP_3_R* and *SERCA* resulted in a reduction in the number of ICTs observed (Movie S13-14). Knocking down gap junctions with *nub*>*Inx2*^RNAi^ had no effect on overall Ca^2+^ activity, but ICTs were primarily smaller spikes (Movie S15). This agrees with Ca^2+^ dynamics observed ex vivo when stimulated by fly extract. Of the three PLC homologs tested, only *nub*>*Plc21C*^RNAi^ influenced the number of ICTs observed (Movie S16-18). RNAi mediated inhibition of Stim and Orai did not influence ICTs in vivo (Movie S19-20).

**Figure 4:**
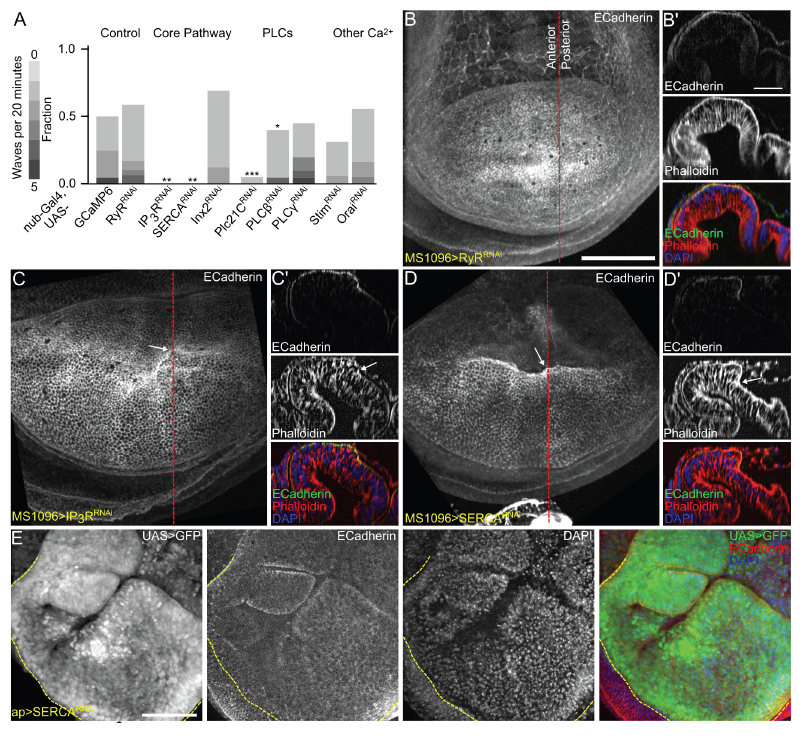
Ca ^2+^ perturbation leads to morphological changes due to perturbed Myosin II. A) ICT activities for calcium pathway knockdowns, PLC homologs, and genes involved in store operated Ca^2+^ signaling (SOCS) was compared to the tester line and an RNAi control for a gene not expressed in the wing disc (*RyR*). ** indicates p < 0.01, *** indicates p < 0.001 by Mann-Whitney *U* test vs. *nub*>*RyR*^RNAi^. B) Z-slice of *MS1096*>*RyR*^RNAi^ disc with Phalloidin staining. Red line indicates position of orthogonal section. B’) Orthogonal section stained with Phalloidin and DAPI. C-D) Maximum z-projection of confocal z-stack of *MS1096*>*IP_3_R*^RNAi^ and *MS1096*>*SERCA*^RNAi^ disc stained for DE-Cadherin respectively. Red line indicates position of orthogonal section. C’-D’) Orthogonal section stained for E-Cadherin, Phalloidin, and DAPI. White arrow indicates position of fold with differential height. E) Disc expressing *GFP* and *SERCA*^RNAi^ driven by *ap-Gal4*, stained for DE-Cadherin and DAPI. Background subtraction was applied to raw stacks using FIJI. Scale bars represent 50 μm.

Previous reports have indicated a role for Ca^2+^ signaling in maintaining wing disc morphology. (Balaji et al., 2017). However, specific connections between levels of Ca^2+^ activity and morphogenetic outcomes remain largely uncharacterized. Differential perturbation of Ca^2+^ signaling components with the Gal4/UAS system provides a powerful means for studying the effect of non-uniform Ca^2+^ signaling on development. We knocked down Ca^2+^ signaling genes in MS1096>RNAi discs and compared the wing phenotypes to the control knockdown of *RyR* (Fig. 4B-D). We observed that *MS1096*>*IP_3_R*^RNAi^ and *MS1096*>*SERCA*^RNAi^ exhibited a decrease in apical area in the dorsal compartment and a specific bend near the boundary of differential perturbations of Ca^2+^ signaling on either side of the D/V axis (Fig. 4C, D), relative to the *MS1096*>*RyR*^RNAi^ control. Additionally, MS1096>*SERCA*^RNAi^ led to ectopic folding throughout the disc near the anterior-posterior compartmental boundary. Furthermore, ap>*SERCA^RNAi^* led to much more severe morphological phenotypes than MS1096>*SERCA*^RNAi^ (Fig. 4E). In older larval discs, *ap*>*SERCA*^RNAi^ showed a significant increase in the size of the dorsal compartment at the expense of the ventral compartment, a near total loss of DE-Cadherin in the dorsal compartment, and ectopic folds throughout the dorsal compartment and along the A/P and D/V boundaries.

### ICTs are stimulated by mechanical stress dissipation

Since tissue mechanics are a crucial factor that regulates the morphology of wing discs and adult wings, the morphological defects caused by perturbed Ca^2+^ signaling suggests that ICTs play a role in defining the mechanical state of the tissue. Additionally, cell-cell connections are crucial for ICTs in the wing disc, because Clone 8 cell culture supports Ca^2+^ oscillations independently of fly extract. Tissue mechanics and cell-cell adhesion are regulated by a group of specialized proteins. For example, the wing disc is tightly anchored (through integrins) to a membrane on the basal side (basement membrane), and has tight cell-cell adhesions (cadherins) along the apical side. The apical side of the wing disc is also under tension due to the contractile cortical actomyosin along the apical surface of each cell (Guillot and Lecuit, 2013). This precisely coordinated balance of forces is necessary for robust development of the wing disc’s size and shape (Ma et al., 2017).

Previous research from our lab has shown that a release of mechanical compression led to ICTs (Narciso et al., 2017), and it suggests that the change of mechanical stress in wing disc is connected to multicellular Ca^2+^ transients. Therefore, we used two methods to perturb the overall mechanical stress in wing discs to further test the hypothesis that mechanical stress dissipation during wing disc development contributes to the rise of ICTs (Fig. 5).

**Figure 5:**
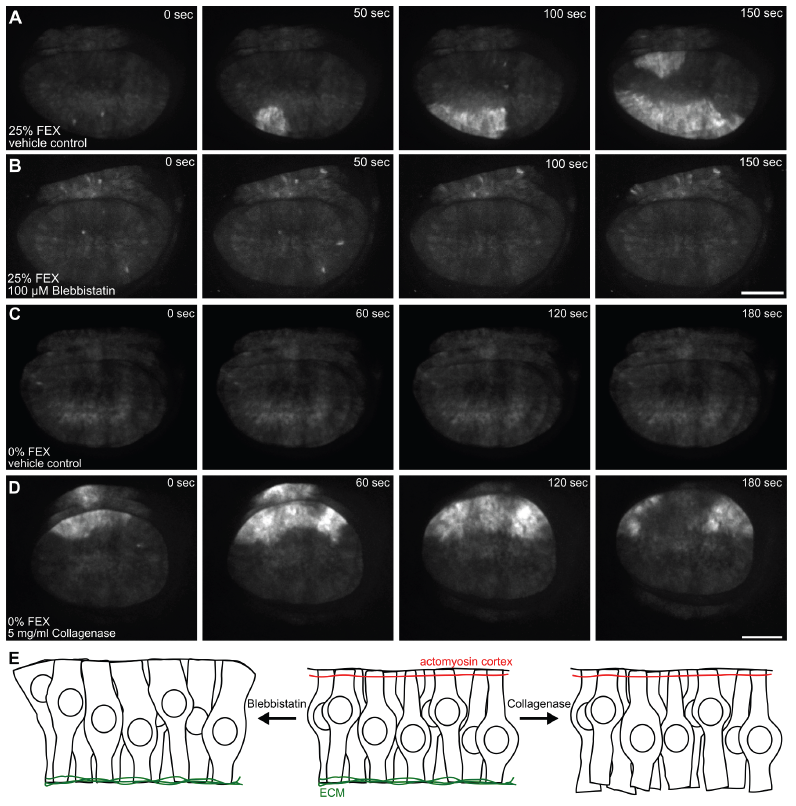
Pharmacological perturbation of tissue mechanics impacts ICTs. A-D) Live imaging of wing discs expressing GCaMP6f under the nubbin driver in ZB media. A) 25% fly extract with 0.6% DMSO, B) 25% fly extract and 100 μM Blebbistatin, C) 0% fly extract with 0.5% PBS, or D) 5 mg/ml collagenase. E) Cartoon representation of the effect of drugs on mechanical state of the tissue. Scale bar represents 100 μm.

First, we used blebbistatin to reduce the overall tension of the wing discs. Blebbistatin inhibits myosin phosphorylation, leading to a decrease in tension both along the apical and lateral surfaces. Ex vivo incubation of wing disc with blebbistatin and 25% fly extract resulted in a loss of ICTs, although Ca^2+^ spikes still occurred (Fig. 5A, B; Movie S21-22), similar to discs cultured in the absence of fly extract (Fig. 1B, 5C), which agrees with previous findings (Balaji et al., 2017).

Second, we used collagenase to quickly release the constraint imposed by the ECM (Fig. 5E). Collagenase breaks down the ECM, and leads to a significant loss of external constraint leading to a flattening and reduction of cell height in the wing disc (Pastor-Pareja and Xu, 2011). Adding collagenase to the ex vivo culture resulted in deformation of the wing disc, and loss of tissue stress. This led to an immediate ICT in all replicates in the chemically defined medium, while no ICTs were observed in any of the vehicle controls (Fig. 5C-D, n = 4; Movie S23-24). Overall, these results demonstrate that dissolution of the ECM stimulates ICTs. This is the first chemically defined perturbation reported for stimulating ICTs in wing discs that does not require wounding cells.

In vivo, the ECM is constantly remodeled, suggesting the natural remodeling of the ECM may explain the stochastic ICTs observed in vivo. Taken together, these findings indicate that different mechanisms of mechanical stress release stimulate ICTs in the wing disc. This also implies that Ca^2+^ mediated stress release is important for ensuring robust wing disc morphogenesis.

## Discussion

In this study, we systematically characterized the morphogenetic roles of the core calcium signaling toolkit by perturbing components of the Ca^2+^ signaling pathway with multiple Gal4 drivers. We identified four classes of Ca^2+^ transients that occur in vivo and ex vivo: spikes, ICTs, ICWs and fluttering discs. We also identified two broad classes of effects on wing morphogenesis: 1) Perturbation of IP_3_ signaling and gap junction communication inhibits ICTs, reduces wing area and perturbs cross veins in adult wings; 2) Inhibition of other channels regulating ER Ca^2+^ homeostasis leads to severe folding defects in adult wings. Further, we inhibited ICTs using blebbistatin and induced ICTs by dissociating the ECM with collagenase, providing significant evidence that release of stress triggers an intercellular Ca^2+^ response.

The finding that fly extract affects ICTs ex vivo in a dose dependent manner indicates that components of fly extract are acting on the disc chemically. Although the identity of the chemical signal remains unknown, it exhibits biochemical activity consistent with being a relatively heat stable protein (Balaji et al., 2017). Moreover, the dependence of ICT activity on developmental stage suggests that ICTs may serve as a readout of some correlated factor during development, such as the growth rate of the tissue, relative morphogen secretion activities or ECM remodeling factors.

Destabilizing the homeostatic concentrations of Ca^2+^ in the ER and the cytoplasm influences Ca^2+^ signaling differently than disrupting transient phenomena such as Ca^2+^ release. SOC channels have been implicated in the control of growth and patterning in wing disc development (Eid et al., 2008; Richard and Hoch, 2015). Interestingly, knockdown of SOC channel components did not strongly impact ICT frequency. However, we cannot rule out that a complete loss of function would be sufficient to block ICTs. The phenotypes of adult wings with knocked down SOC channel components resembled wings expressing *SERCA*^RNAi^. These stronger phenotypes are likely due to additional requirements of ER Ca^2+^ for proper processing of Notch and other morphogens (Ilagan and Kopan, 2013).

Defects in the cross veins suggests that morphogen patterning in the wing is perturbed when Ca^2+^ signaling is inhibited (O’Connor et al., 2006; Yan et al., 2009; Yu et al., 1996). We show that knockdown of IP_3_R, Inx2 and Plc21C results in partial loss of the cross veins. It suggests that a decrease of Ca^2+^ signaling leads to decrease of Dpp signaling, because late pupal veins secrete Dpp. We also demonstrate that knockdown of Inx2 results in smaller wings, which suggests a decreased level of Dpp signaling, since a major role of Dpp signaling in the wing disc is the control of tissue size (Restrepo et al., 2014; Richard and Hoch, 2015). A recent study of a potassium channel *irk2* also shows the reduction in Ca^2+^ signaling correlates with the reduction in the rate of Dpp release (Dahal et al., 2017). Taken together, the evidence is consistent with coupling of Ca^2+^ dynamics and Dpp signaling (Dahal et al., 2017). Other pathways, such as Wg, Notch and EGF pattern veins as well (Crozatier et al., 2004; Werner et al., 2010). Ca^2+^ signaling also may interact with major morphogen pathways and plays a major role in fine-tuning high levels of morphogen signals. In an accompanying study (Brodskiy et al., DEVELOP/2017/159269), we test and confirm a hypothesis that time-averaged statistics in Ca^2+^ signaling dynamics are downstream of morphogenetic signaling. Through perturbations to Hh-signaling, we further confirm that these Ca^2+^ signaling signatures reflect integration by Ca^2+^ signaling activity.

The characteristic phenotype of bent wings, caused by the knockdown of IP_3_R, Inx2 and Plc21C, provide additional clues to the potential downstream targets of Ca^2+^ signaling involved in morphogenesis. Examples include *Curly* and *Wavy*, which both exhibit a bent wing phenotype. The *Curly* mutation has been traced to NADPH dual oxidase (*Duox*), which generates reactive oxygen species (Hurd et al., 2015). This phenotypic similarity suggest that Ca^2+^ signaling may regulate the generation of reactive oxygen species in wing discs (Razzell et al., 2013). *Wavy* is a negative form of IP_3_K2 that generates IP_4_ from IP_3_ (Dean et al., 2015), which causes an increase in IP_3_ concentration. An increase in IP_3_ concentration can lead to an overstimulated phenotype with high levels of Ca^2+^ signaling that prevent ICTs. These potential mechanisms by which perturbation of Ca^2+^ signaling results in the disrupted epithelial folding and bent wings remain to be explored.

We hypothesize that ICTs dissipate mechanical stress during in vivo growth of tissues. Previously, we showed that when mechanical loading on a wing disc was released, a long-range ICW is stimulated (Narciso et al., 2017). Here, we demonstrate that the dissolution of the ECM by collagenase induces ICTs and that ICTs occur naturally in vivo. Both the abovementioned processes involve the remodeling of the ECM, which accompanies the buildup and release of mechanical tension. In early stage *Drosophila* embryos, Ca^2+^ spikes in single cells, but ICTs have not been reported (Markova et al., 2015). This can be explained by a lack of organismal size changes during embryo development or ECM remodeling during early embryonic stages (Lye and Sanson, 2011). As a serum containing many proteases, fly extract may stimulate ICTs through the disruption of ECM by proteases or other chemical signaling factors. Moreover, the dependence of ICT activity on developmental stage may also be explained by the reduction of the rate of ECM remodeling that must occur as discs reach terminal sizes. These observations suggest that ICTs may be a natural consequence of ECM remodeling during tissue growth, and may serve as an important feedback mechanism for controlling tissue shape. Thus, Ca^2+^ signaling modulates morphogen signaling and tissue mechanics, which warrants future investigations.

## Materials and Methods

### Fly culture

Gene perturbations to the core Ca^2+^ signaling pathway (*Gene X*^RNAi^ or *Gene X* overexpression) were crossed to a genetic tester line (*nub-GAL4*, *UAS-GCaMP6f*/*CyO*) that enables visualization of Ca^2+^ signaling with down regulation of genes in the wing disc in vivo. For imaging of larval and adult morphology, two genetic tester lines were used (ap-Gal4 (Milán et al., 1997), and MS1096-Gal4 (BL#8860). When possible, multiple independent RNAi lines were tested for each gene investigated (Table S1). Stocks are obtained from Bloomington *Drosophila* Stock Center as indicated by stock number (BL#). Progeny wing phenotypes are from F1 male progeny emerging from the nub-Gal4, UAS-GCaMP6f/CyO x UAS-X cross. The tester line (w1118; nubbin-GAL4, UAS-GCaMP6f/CyO) was generated by recombining P{UAS-Dcr-2.D}1, w1118; P{GawB}nubbin-AC-62 (BL#25754) with w1118; P{20XUAS-IVS-GCaMP6f}attP40 (BL#42747). Flies were raised at 25**°**C and 12-hour light cycle.

### Adult wing imaging

Whole flies and intact wings were imaged at 1.6x using an Olympus SZX7 dissection microscope (Waltham, MA). To obtain high-resolution images of vein patterning phenotypes, wings were detached and mounted in Euparal and imaged at 4x on EVOS™ FL Auto Imaging System (Thermo Fisher, Waltham MA). For larger wings, images were digitally stitched and rotated with FIJI (Schindelin et al., 2012).

### In vivo larval wing disc imaging

Wandering 3^rd^ instar *nub-GAL4*, *UAS-GCaMP6f*/*CyO* larvae were collected for imaging and rinsed in deionized water. They were dried and then adhered to a coverslip for imaging with scotch tape covering the larvae. The larvae were attached with their spiracles facing toward the coverslip to ensure the wing discs were aligned toward the microscope. The EVOS™ FL Auto Imaging System was used to image the larvae. The larvae were imaged at 20x magnification for 20 minutes; images were taken every 15 seconds.

### Wing disc ex vivo imaging

Wing discs were dissected, cultured, and imaged as described in (Zartman et al., 2013) from 3^rd^ instar larvae and cultured and imaged in the organ culture media on a cover slip based on our previously developed protocol. Each disc was immobilized by either a cell culture insert (EDM Millipore) with truncating the legs, or by culturing in a REM chip device (described in (Narciso et al., 2017)). The setup was then transferred to a confocal microscope for Ca^2+^ activity imaging. Imaging was performed on a Nikon Eclipse T*i* confocal microscope (Nikon Instruments, Melville, NY) with a Yokogawa spinning disc and MicroPoint laser ablation system (Andor Technology, South Windsor, CT). Image data were collected on an iXonEM+ cooled CCD camera (Andor Technology, South Windsor, CT) using MetaMorph^®^ v7.7.9 software (Molecular Devices, Sunnyvale, CA). All experiments were performed within 20 minutes of dissection to minimize time in culture. Discs were imaged at a three z-planes with a step size of 10 μm, 20x magnification and 10-second intervals for a total period of one hour, with 200 ms exposure time and a 50mW 488 laser (Andor Technology, Belfast, UK) at 50% intensity. Microscopy resulted in 4D time-lapse data (512 pixels by 512 pixels by 3 z-planes by 361 time points). The z-stack data was max-projected in FIJI (Schindelin et al., 2012) to yield z-projected time-lapse videos.

### Wing disc immunohistochemistry

Discs were fixed after dissection by immersing in 4°C 10% normal buffered formalin for 15 mins. Discs went through 3X quick rinses and 3X 10-min rinses with PBT in room temperature. Remove the last rinse, add blocking solution (5% goat serum), and block for 30 minutes at room temperature. Remove block and replace with primary antibody solution. Stain with primary antibodies for 5 hours at room temperature or overnight at 4C. Remove the primary antibodies and conduct 3X quick rinses and 3X 10-min rinses with PBT. Remove the last rinse and add secondary antibody solution, which includes secondary antibodies diluted in blocking solution. Stain with secondary antibodies for 2 hours at room temperature. The secondary solution is then removed and discarded. Three quick rinses and three 10-min rinses with PBT were performed. Discs were mounted on thin coverslips in VECTASHIELD (H-1000, Burlingame, CA), sealed with nail polish. Antibody sources and concentrations are described in Table S2.

### In vivo calcium signaling imaging analysis

For each video, the total number of distinct initiation sites of Ca^2+^ ICTs and ICWs was measured. Videos were assigned random names and were analyzed in a random order. The total number of events in 20 minutes was reported. Separately, each movie was divided into 135 second clips and each clip was classified as having either no activity, spikes, ICTs, ICWs, fluttering, or being unanalyzable because of larval motion. This was done with a custom MATLAB script which divided each movie into clips. One random clip was looped at a time until all the data had been classified. Afterwards, the fraction of time spent in each regime was reported for each condition.

### Wing vein phenotype analysis

Wing images were randomly renamed and assessed in a random order for ACV and PCV phenotypes. Whole wings were segmented and wing area was measured using FIJI (Schindelin et al., 2012).

### Generation of stable Clone 8 GCaMP6f line

Ca^2+^ sensors for Clone 8 cell transfections were made from the pGP-CMV-GCaMP6f plasmid (Addgene #40755) as described in SI Materials and Methods. The fast calcium sensor ORF was then cloned into the vector pAc5-STABLE1-blast (a gift from Dr. J. D. Sutherland, (González et al., 2011)) for insect cell expression and selection of stable clones using blasticidin.

### Insect cell microscopy

Stably-transfected Clone 8 cells were conditioned to the chemically defined ZB media for at least two passages before performing the experiments. Cultures were kept with the selection antibiotic Blasticidin at 25 μg/mL. Cells were treated with Deep Red CellTracker 630/650 dye for 30 minutes (Thermo Fisher Scientific Inc., 2014). 50,000 cells per well were plated on 96 well plates (Greiner CELLSTAR®) in 100 μl of ZB media and allowed to adhere for 3 hours at 25°C, and sealed with Parafilm (Bermis, Neenah, WI). One site in each of 60 wells was selected and GCaMP6 was imaged for 9 minutes at 1 second intervals with a 488 laser, 300 gain, and 200 ms exposure time at a Nikon CFI Plan Apo 20x/0.75 DIC N2 objective. In between 9-minute imaging sessions, each site was imaged with the 488 and 640 lasers to obtain long-term time-lapse video for each site. Long-term imaging data revealed that cells continued to divide and migrate throughout the 12-hour imaging time. ROIs were generated and intensity profiles were collected using FIJI (Schindelin et al., 2012).

### Statistical analysis

The Mann–Whitney *U* test was used to compare number of in vivo Ca^2+^ events between each UAS cross and the GAL4 tester line control because data followed a power law distribution and thus was not normally distributed. Standard hypothesis testing could not be carried out due to the high number of UAS cross variants being compared to the GAL4 control. the family-wise error rate is the probability of making a false discovery. The family-wise error rate increases as the number of hypotheses tests being performed increases. Multiplicity corrections were applied to the p-values obtained from the Mann–Whitney *U* test via the Benjamini–Hochberg procedure to avoid significant false discoveries (Benjamini and Hochberg, 1995). Utilizing this correction procedure allowed for controlling the false discovery rate at a significance level of α = 0.05. Significance of developmental stage trend was determined by t-test comparing a linear regression model to null model.

### Image processing

For time-lapse data relevant images were selected from time stacks using FIJI (Schindelin et al., 2012). Brightness and contrast was consistent for each frame within each figure and was set with a MATLAB function. Immunohistochemistry data was processed using FIJI (Schindelin et al., 2012). Background subtraction with pixel radii of 10 (DAPI and Phalloidin) and 20 (GFP and ECadherin) was applied to each stack using the sliding paraboloid setting in FIJI. Maximum z projections were generated using FIJI. Where possible, the peripodial membrane was digitally removed by removing the relevant z-slices prior to z projection. Stacks were then digitally rotated such that the posterior compartment was on the right, and flipped such that the dorsal compartment was on the top of the image. Stacks were cropped to 512 x 512 pixels to maintain image sizes consistent with the camera sensor. Orthogonal projections were obtained using the Reslice function in FIJI. Additional background subtraction with pixel radii of 5 (Phalloidin and ECadherin) and 1 (DAPI) was applied to each orthogonal projection using the sliding paraboloid setting in FIJI. Orthogonal projections were then cropped such that images generated from different stacks were identical in size. Cropped orthogonal projections were then flipped such that the ventral compartment was on the left.

## Acknowledgements

This work was supported in part by NSF Awards CBET-1403887 and CBET-1553826, NIH Grant R35GM124935, ND Advanced Diagnostics and Therapeutics Discovery Fund and Harper Cancer Research Institute Research like a Champion awards (QW), Walther Cancer Foundation Interdisciplinary Interface Training Project (PB), and the Notre Dame Advanced Diagnostics & Therapeutics Berry Fellowship (CN). The authors gratefully acknowledge the Notre Dame Integrated Imaging Facility. The authors would like to thank Kara Synder, Ninfamaria Arredondo-Walsh and Roy Brooks for technical assistance, the Bloomington Stock Center for fly stocks, and the Developmental Studies Hybridoma Bank for antibodies.

## Extended Data

**Fig. S1.**
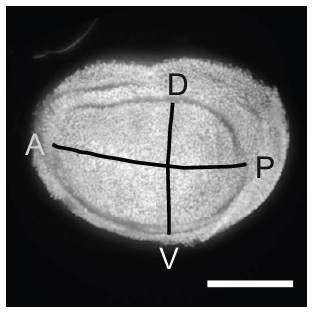
*nubbin-Gal4* driven expression is uniform in the wing disc pouch. *nub-Gal4*>*UAS-RFP* is expressed in the pouch. The intensity of RFP is uniform throughout the disc, suggesting uniform expression of GCaMP6 calcium sensor. A: anterior; P: posterior; D: dorsal; V: ventral. Scale bar: 50 *μ*m.

**Fig. S2.**
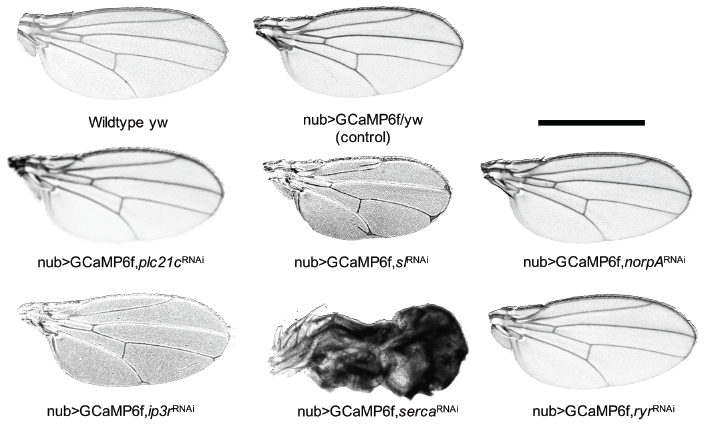
Example wing phenotypes from Ca ^2+^ genes perturbation with nub-Gal4 driver. Shown here are representative images from each genetic condition. *sl^RNAi^* led to smaller wings, and the end of the veins bifurcated. *norpA^RNAi^* and *ryr^RNAi^* did not change wing morphology. *ip3r^RNAi^* resulted in the bifurcation of L4 & L5 veins. Control was generated from crossing the tester line (nub>GCaMP6/CyO) to yw. Scale bar: 1 mm.

**Fig. S3:**
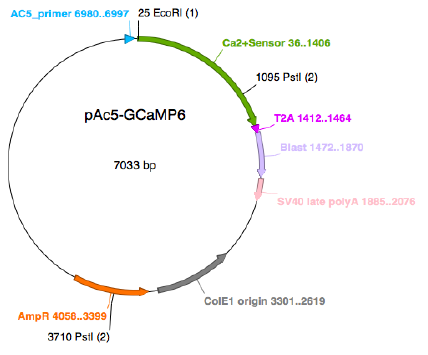
Schematic of insect cell calcium sensor plasmid pAc5-GCaMP6f.

## SI Materials and methods

### Generation of stable Clone 8 GCaMP6f line

A double digestion of pGP-CMV-GCaMP6f (Addgene #40755) with enzymes NotI and BglII generated the target fragment (1,352 bp) which was purified from 1% low gelling temperature Agarose gel (Sigma A9414) using QIAquick gel extraction kit (Qiagen 28704) and treated with T4 DNA Polymerase (Promega M421A) to create blunt ends. The target was then cloned into the vector pAc5-STABLE1-blast (Addgene Ac5-STABLE1-neo plasmid # 32425 modified version containing Blasticidin instead of Neomycin, provided as a gift from Dr. J. D. Sutherland) for insect cell expression. Briefly, the vector was first treated with XbaI and HindIII for a double digestion to remove original EGFP module. The modified vector, 5,658 bp, was gel purified and blunted as described above. Next the treated vector was ethanol precipitated and dephosphorylated with Thermosensitive Alkaline Phosphatase (Promega M9910). The insert was ligated into the modified vector using T4 DNA ligase (Promega, M8221) with a 2:1 molar ratio insert: vector and overnight incubation at 16 °C. Ligation products were transformed into DH5α competent cells (Promega, L122a) and screened with PstI for the correct orientation of insert. The final insect cell Ca2+ sensor expression plasmid was named pAc5-GCaMP6f and it is shown in (Fig. S3).

Drosophila Clone 8 (Clone 8) cell line (CME w1 Clone 8+ Melanogaster, dorsal mesothoracic disc, modENCODE line #151) was obtained from the DGRC and maintained in Clone 8 media in a humidified incubator at 25°C in plug seal tissue culture flasks (Clone 8 media: M3 + 2% FCS + 5 μg/mL human insulin + 2.5% fly extract). Clone 8 cells were transfected with the Trans IT system (Mirus) and selected in Blasticidin at 25 ug/mL in Cl. 8 media. Stable clones were obtained after two months of antibiotic selection in well plates.

**Table S1.**
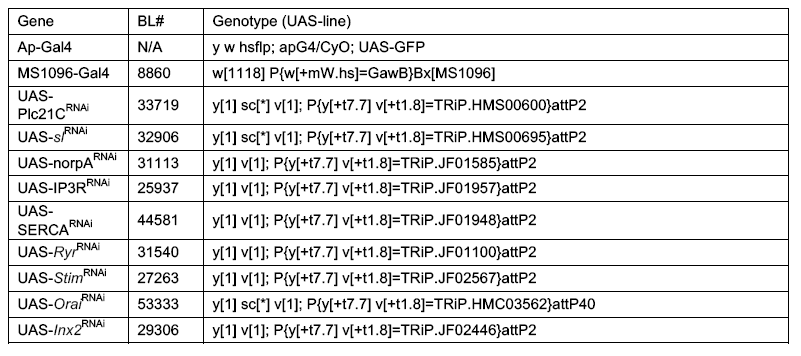
Drosophila lines.

**Table S2.**
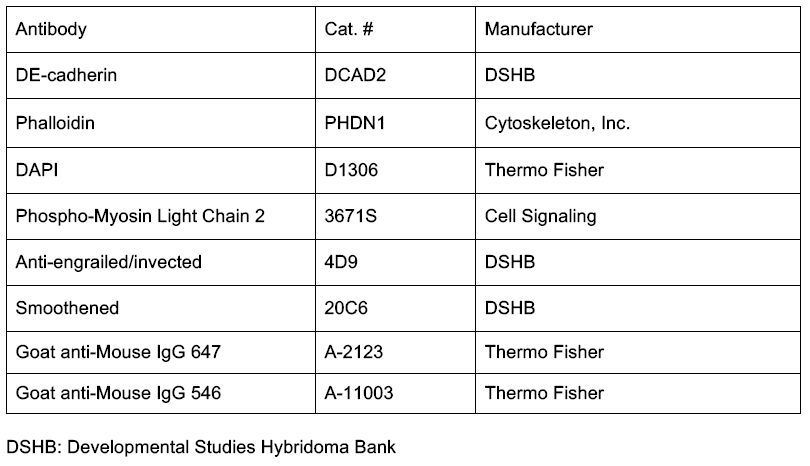
Antibody sources.

**Table S3.**
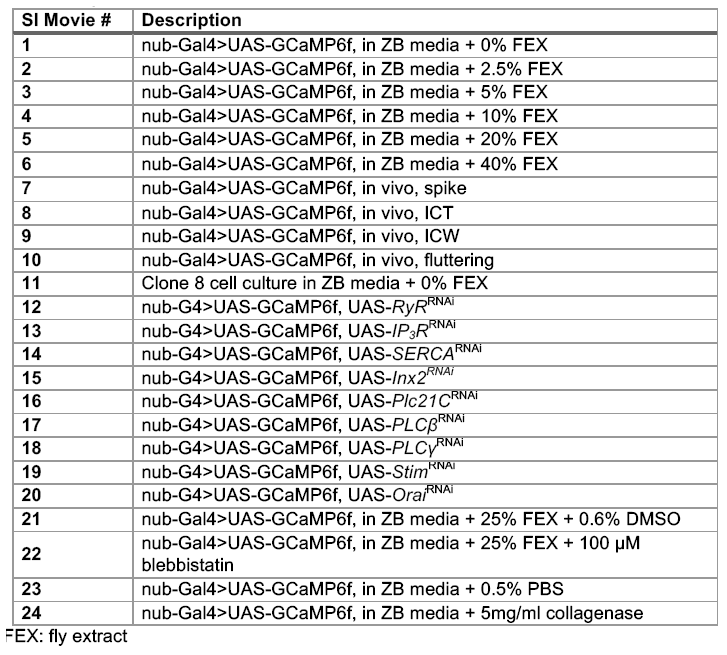
Extended data movies.

